# Repeated administration of cannabidiol decreases splenic lymphocyte numbers in rats: involvement of CB_2_ receptors

**DOI:** 10.1101/2025.02.24.639985

**Authors:** Tara H. Turkki, Maciej. M. Jankowski, Wojciech Glac, Piotr Badtke, Viviane M. Saito, Artur H. Swiergiel, Bogna M. Ignatowska-Jankowska

## Abstract

Cannabidiol (CBD), the major non-psychotropic compound of *Cannabis sp.*, is an effective treatment for inflammatory and autoimmune diseases and produces various anti-tumor effects but the mechanisms of its long-term actions *in vivo* remain unclear. We have previously shown that CBD administration (5 mg/kg) in healthy rats significantly decreased lymphocyte numbers in peripheral blood, involving B, T CD4+ and T CD8+ lymphocyte subsets, but not natural killer (NK) cells. To examine the effects of CBD on lymphocyte subsets in the spleen and NK cellular cytotoxicity (NKCC), adult male Wistar rats (n = 63) were administered intraperitoneal injections of CBD (2.5 or 5 mg/kg/day) for 14 consecutive days and lymphocyte counts were obtained using flow cytometry. NKCC in the peripheral blood and spleen was quantified using a Chromium-51 release assay. Furthermore, CB_2_ receptors were blocked using selective receptor antagonist AM630 (1 mg/kg). The results indicate that repeated administration of CBD at a dose of 5 mg/kg/day resulted in a decrease in splenic lymphocyte number, involving T and B lymphocytes but not NK cells. The decrease in lymphocyte number was partially blocked by pretreatment with CB_2_ receptor antagonist while no changes in NKCC were observed following CBD administration. These results reveal that in healthy rats, CBD produces similar lymphopenic effects in the spleen as it does in peripheral blood and that the effects of CBD on lymphocyte numbers *in vivo* are at least partially mediated by CB_2_ receptors.

## Introduction

Cannabidiol (CBD) is one of the major non-psychotropic phytocannabinoids found in *Cannabis sp*. The plant has been used in medicine throughout history, but psychotropic side effects have largely limited its use in therapy. Unlike the major psychotropic compound of cannabis, THC (1′9-tetrahydrocannabinol), CBD displays a low affinity to cannabinoid CB_1_ and CB_2_ receptors (Howlett et al., 2002; Pertwee, 2008). However, CBD can inhibit the effects of CB_1_ and CB_2_ cannabinoid receptor agonists and was shown to behave as a CB_2_ receptor inverse agonist *in vitro* (Pertwee, 2008; Thomas et al., 2007) as well as a negative allosteric modulator of CB_1_ (Laprairie et al., 2015). Moreover, CBD is known to affect cannabinoid receptors indirectly by modulation of alternative pathways such as endocannabinoid signaling, for example by inhibiting FAAH (Bisogno et al., 2001; McPartland et al., 2015). Much of the therapeutic use of cannabinoids is focused on those devoid of psychotropic properties, namely CBD as it is also most abundant in hemp plants (Izzo et al., 2009; Mechoulam et al., 2007; Pisanti et al., 2017). Like many other cannabinoids, CBD produces analgetic, antiemetic, neuroprotective, and immunomodulatory effects (Izzo et al., 2009; Mechoulam et al., 2007) and because of low toxicity and safety of use in humans (Cunha et al., 1980; Rosenkrantz et al., 1981; Zuardi, 2008), it has been proven to have wide therapeutic applications in the treatment of various diseases (Nichols and Kaplan, 2020; Pisanti et al., 2017; Zuardi, 2008).

CBD has been shown as effective in animal models of autoimmune diseases such as rheumatoid arthritis (RA), type I diabetes, and multiple sclerosis (MS) via its anti-inflammatory and anti-autoimmune effects (Kozela et al., 2011; Malfait et al., 2000; Weiss et al., 2008). It has also been shown to alleviate neuropathic pain both in animal models and humans (Costa et al., 2007, 2007; Gregorio et al., 2018; Nichols and Kaplan, 2020; Russo et al., 2016; Urits et al., 2020). Moreover, CBD is known to produce antitumor effects decreasing the proliferation and migration of various tumor cell lines *in vitro*, as well as tumor growth and metastasis *in vivo* (Ligresti et al., 2006; Massi et al., 2004). While the antitumor effects of CBD have been demonstrated extensively, its biological effects on different human cancers are dependent on cell type and origin (Valenti et al., 2022). Other shown benefits of CBD specifically are improved sleep and mood quality (Chagas et al., 2014, 2013; Notcutt et al., 2004; Serpell et al., 2014), as well as anxiolytic and antipsychotic effects (Crippa et al., 2018; Guimarães et al., 1990; Zuardi, 2008). Most recently, CBD (Epidiolex), has been FDA-approved for the treatment of drug-resistant or refractory epilepsy (Sekar and Pack, 2019).

The mechanisms of CBD’s influence on the immune system have been thoroughly investigated *in vitro*, where it was found to induce apoptosis, decrease cell proliferation, cytokine production, and induction of regulatory cells. Moreover, *in vivo* effects of CBD have been shown to be mostly immunosuppressive and anti-inflammatory (Atalay et al., 2020; Nichols and Kaplan, 2020). However, the exact mechanisms through which CBD affects lymphocyte number *in vivo* and whether *in vitro* observations are relevant to physiological conditions (Lee et al., 2008; Wu et al., 2008) is unclear. There is contradictory data concerning the involvement of CB_2_ receptors in the effects of CBD on the immune system but some studies indicate that CBD may have the ability to exert its effect *via* CB_2_ receptors (Castillo et al., 2010; Ignatowska-Jankowska et al., 2011; Sacerdote et al., 2005).

Despite the wide therapeutic potential of CBD (Izzo et al., 2009; Mechoulam et al., 2007) in the treatment of inflammatory and autoimmune diseases, the effects of prolonged, systemic administration of CBD on the immune system and its mechanisms of action remain unclear. Our previous studies revealed a significant decrease in lymphocyte number in the peripheral blood of rats following systemic CBD administration for 14 consecutive days (at a dose of 5 mg/kg that has been found to be most effective in animal models of inflammatory and autoimmune diseases). The present study aimed to examine effects of repeated administration of CBD on the lymphocyte numbers (B, T, NK and T CD4+ and T CD8+ subsets) in the spleen of rats. Furthermore, the study aimed to determine whether effects of administration of CBD on lymphocyte numbers are mediated by CB_2_ cannabinoid receptors that have an important role in regulation of immune response by cannabinoids. Moreover, because our previous studies indicated that CBD administration may increase numbers of NK cells that are responsible for primary antitumor and antiviral innate immune response, this study aimed to examine whether repeated treatment with CBD affects cytotoxic activity of NK cells against tumor cells.

## Materials and Methods

### Animals

The subjects (n=63 (36 + 27)) were adult male Wistar rats (Experiment 1: R. Grabowski Laboratory Animals Breeding, Gdansk; Experiment 2: Trojmiejska Akademicka Zwierzetarnia Doswiadczalna, Gdansk, Poland), 10 weeks old, 250 ± 20 g at the beginning of experiments. Animals were housed in cages of 4, with free access to food and water. The animal room was maintained at a temperature of 21 ± 1°C with humidity 55 ± 10 %, under 12-h light cycle (lights on from 6:00 to 18:00). For 14 days prior to the start of experiments rats were handled daily and adapted to the presence of the experimenter to minimalize stress evoked by experimental procedures.

The principles for the care and use of laboratory animals in research, as outlined by the Local Ethical Committee (permission number: 33/2008), were strictly followed and all the protocols were reviewed and approved by the Committee. All efforts were made to minimize animal discomfort, and the number of animals used.

For experiment 1, animals were divided randomly into three groups, and for 14 consecutive days received intraperitoneal injections of CBD at a dose of 2.5 mg/kg/day (Group I, n=9), 5 mg/kg/day (Group II, n=9), day or the vehicle (Group III, n=9). For experiment 2, animals were divided randomly into four groups and received two intraperitoneal injections for 14 consecutive days. The experimental group was pre-treated with CB_2_ receptor antagonist AM630 at a dose of 1 mg/kg/day followed by injections of CBD at a dose of 5 mg/kg/day (Group I, n=9). The three control groups were pre-treated with the vehicle, followed by 5 mg/kg/day of CBD (Group II, n=8), pre-treated with 1 mg/kg/day of AM630 followed by vehicle (Group III, n=9), or received vehicle for both treatments (Group IV, n=11).

### Drug administration and experimental design

CBD (*Cannabidiol*, Lipomed, Arlesheim, Switzerland) and AM630 (6-iodo-2-metyl-1-[2-(4-morpholinyl)etyl]-1H-indol-3-yl](4methoxyphenyl)methanone; Tocris Bioscience, Bristol, United Kingdom) were dissolved in a vehicle solution containing Cremophor (Sigma Aldrich, Germany), 99.9% ethanol, and saline (0.9% NaCl) in a 1:1:18 ratio, which was also used in control groups. For 14 consecutive days animals received intraperitoneal injections of vehicle or CBD at doses of 2.5 or 5 mg/kg/day. Doses were based on recent literature, the effects of CBD and AM630 in animal models, and our preliminary experiments. AM630, a selective antagonist of CB_2_ cannabinoid receptors was administered intraperitoneally at dose of 1 mg/kg, 15 min prior to injection of CBD or vehicle (Bisogno et al., 2009, 2001; Howlett et al., 2002; Ibrahim et al., 2003; Malan et al., 2001). All solutions were administered at fixed time points for each animal in a volume of 1 ml/kg and were prepared immediately before use.

### Blood and Spleen Sampling

Blood samples (3 mL) were drawn to measure the cytotoxic activity of NK cells on experimental day 14, 60 minutes after the last CBD or vehicle injection. Samples were collected by cardiac puncture under halothane anesthesia (*Narkotane* Zentiva, Prague, Czech Republic) according to the standard procedure used in our laboratory (Wrona et al., 2004). Blood sampling was limited to 2 minutes after taking the animal from the home cage, to avoid effects of stress on immune parameters. Immediately after blood sampling, the rats were infused with a lethal dose of intracardial pentobarbital (450 mg/kg, ∼160 mg per animal, Morbital, Biowet, Pulawy, Poland) and their spleens were harvested. Spleen weight and volume were recorded immediately followed by homogenization, filtration by pushing through nylon mesh bags, and suspension in Hanks’ Balanced Salt Solution (HBSS, Sigma-Aldrich) with sodium bicarbonate (Merck, USA) for further analysis.

### Analysis of leukocyte and lymphocyte subset number

In spleen samples, distribution of lymphocyte subsets was determined by flow cytometry using a three-color immunofluorescent antibody staining procedure for determination of T (CD3+), B (CD45ra+), NK (CD3-CD161a), as well as T helper (CD3+CD4+) and T cytotoxic (CD3+CD8+) lymphocyte subsets (CD3-FITC/CD45RA-PC7/CD161A-APC and CD3-FITC/CD4-PC7/CD8-APC antibodies, IO Test kit, Beckman Coulter, Immunotech, Marseille, France) as described previously (Ignatowska-Jankowska et al., 2009; Jankowski et al., 2023). All antibodies were added and incubated with whole blood according to the manufacturer’s instructions. Erythrocytes were lysed (1 mL Versalyse, Beckman Coulter) and the lymphocytes fixed (25 μL Fixative Solution, Beckman Coulter). Samples were analyzed within 2 hours, using a FC500 flow cytometer (Beckman Coulter). Total leukocyte number was assessed by hematocytometer (Neubauer chamber). Percentage numbers of leukocyte populations in blood and spleen: lymphocytes, neutrophils, eosinophils, and monocytes were determined by morphological methods (May-Grünwald and Giemsa staining). The total number of each leukocyte subset was calculated by multiplying the total leukocyte number and the percentage of the individual subset.

### NKCC assay

Cytotoxic activity of NK cells in peripheral blood and spleen was quantified using a chromium-51 release assay. As target cells YAC-1 murine lymphoma cell line was used. Washed in complete medium target cells (5 x 10^6^) were labeled with 100 μCi of Na_2_^51^CrO_4_ (Radio Chemical Center, Otwock – Swierk, Poland) at 37°C for 1h. Labeled target cells were washed with RPMI containing 2% FCS and adjusted to 1 x 10^5^ / ml in complete medium. Target cells were cultured in round-bottomed micro-well plates (Nunc, Roskilde, Denmark) with concentration of effector cells E: T = 50:1 (in a total volume of 200 μl) (Exp) in triplicate under standard culture conditions for 4 h. Spontaneous chromium-51 release wells (Sp) had target cells plus 100 μl of complete medium and the maximum release wells (Max) contained target cells plus 100 μl of complete medium with 5 % Triton X-100 (Serva Feinbiochemia, Heidelberg / New York). The assay was terminated at 4 h by centrifuging the plates. Subsequently, 100 μl of supernatant was removed from each well, and triplicate samples were counted in gamma-counter (Wizard 1470) for 1 min. Percentage of the cytotoxicity was calculated as [(Exp – Sp) / (Max – Sp)] x 100. Results of NKCC assay are presented as lytic units, LU_20_ which were calculated as E_std_ / E:T_20_ x T_std_, where E_std_ states for the standard number of effector cells (10^7^), T_std_ states for the standard number of target cells (2 × 10^4^), and E:T_20_ effector-to-target cell ratio, where lysis of 20% of target cells occurs.

### Corticosterone concentration measurements

Blood samples (0.4 mL, with EDTA) were centrifuged for plasma sample collection and storage at −70 °C until use. Quantitative determination of corticosterone concentration in rat plasma was performed using a commercial radioimmunoassay kit (ICN Biochemicals, Inc., Costa Mesa, CA, USA) and Wizard 1470 gamma counter (Pharmacia LKB, Turku, Finland). The assay was performed according to the manufacturer’s instructions.

### Statistical analysis

Data were analyzed with one-way or two-way analysis of variance (ANOVA) followed by Tukey’s or Dunnett’s *post hoc* tests as appropriate. Results are presented as Mean ± SEM and statistical significance threshold is p<0.05.

## Results

### Experiment 1

Total splenic leukocyte (splenocyte) number (including 97-98 % lymphocytes, 1.3-2.3 % neutrophils, <1 % monocytes, and eosinophils) in rats treated with CBD at a dose of 5 mg/kg decreased by 23.8 % as compared to vehicle (1.189 ± 0.087 x 10^6^ in vehicle, 1.188 ± 0.131 x 10^6^ in 2.5 mg/kg, 0.906 ± 0.062 x 10^6^ in 5 mg/kg groups), but this effect was not significant (One-way ANOVA, Dunnett’s post hoc test, p = 0.058, F_2.22_ = 3.25, Fig. 1A). Similarly, there was no significant effect on total splenic lymphocyte number (Supp. Fig 2).

**Fig. 1.**
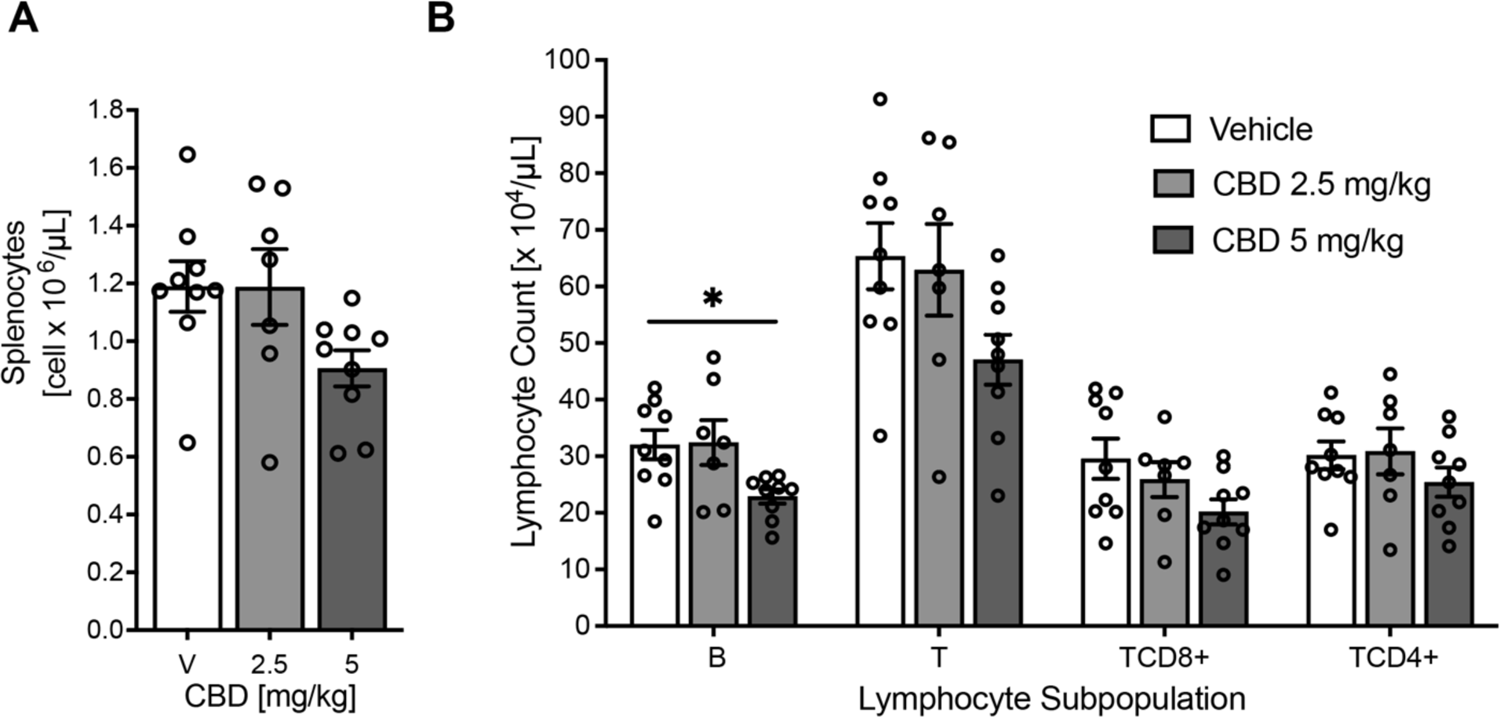
Decreased lymphocyte numbers in the spleen following administration of cannabidiol for 14 consecutive days. (A) Total number of splenocytes. (B) Total numbers of splenic lymphocyte subsets: B cells, T cells, T CD8+, and T CD4+. Data presented as means ± SEM, n = 7-9 per group. One-way ANOVA followed by Dunnett’s post hoc test, *p < 0.05. CBD: cannabidiol, V: vehicle.

Within the splenic lymphocyte subpopulations repeated administration of CBD at a dose of 5 mg/kg resulted in a significant 28.5 % decrease in total B lymphocyte number in the spleen as compared to the vehicle-treated group (p = 0.025, F_2.22_ = 4.36). This was a decrease from 32 ± 0.2 x 10^4^ cells/μL in the vehicle-treated group to 22.9 ± 1.2 x 10^4^ cells/μL in the group receiving 5 mg/kg of CBD (Fig. 1B). A dose of 2.5 mg/kg did not change B cell number (32.4 ± 2 x 10^4^ cells/μL) or any other of the assessed splenic lymphocyte subset numbers (Fig. 1B). The change in total T cell number was not significant (One-way ANOVA, Dunnett’s post hoc test, p = 0.078, F_2.22_ = 2.88, Fig1B). This slight decrease in T cell number was mainly due to a decrease in T CD8+ lymphocyte number by 31.8 % compared to the vehicle (One-way ANOVA, Dunnett’s post hoc test, p = 0.094, F_2.22_ = 2.65) but also T CD4+ lymphocyte number decreased by 15.8 % compared to the vehicle (One-way ANOVA, Dunnett’s post hoc test, p = 0.37, F_2.22_ = 1.03, Fig. 1B). However, these effects were not significant. Overall, no significant change in the percentage distribution of B, T, T CD8+ and T CD4+ cells in lymphocyte population was observed (not shown), suggesting a proportional decrease in the numbers of B and T cell subsets.

Furthermore, treatment at either dose of CBD caused no significant changes in counts or percentages of total splenic lymphocytes, as well as granulocytes or monocytes (Supp. Fig. 2-3). Additionally, neither dose induced significant changes in granulocyte counts or percentages, as well as total lymphocyte numbers in the spleen. The same trends were observed in the blood (Supp. Fig. 2-3).

CBD at either dose did not affect total or percentage splenic NK cell numbers (p = 0.324, F_2.22_ =1.19) or fraction of total lymphocytes (p = 0.790, F_2,22_ = 0.24) (Fig. 2A-B). Furthermore, no significant changes in NK cytotoxic activity in the spleen or peripheral blood were observed (p=0.464, F_2.24_ = 0.79, p = 0.154 and F_2.39_ = 1.962, in the spleen and blood respectively, Fig. 2C-D).

**Fig. 2.**
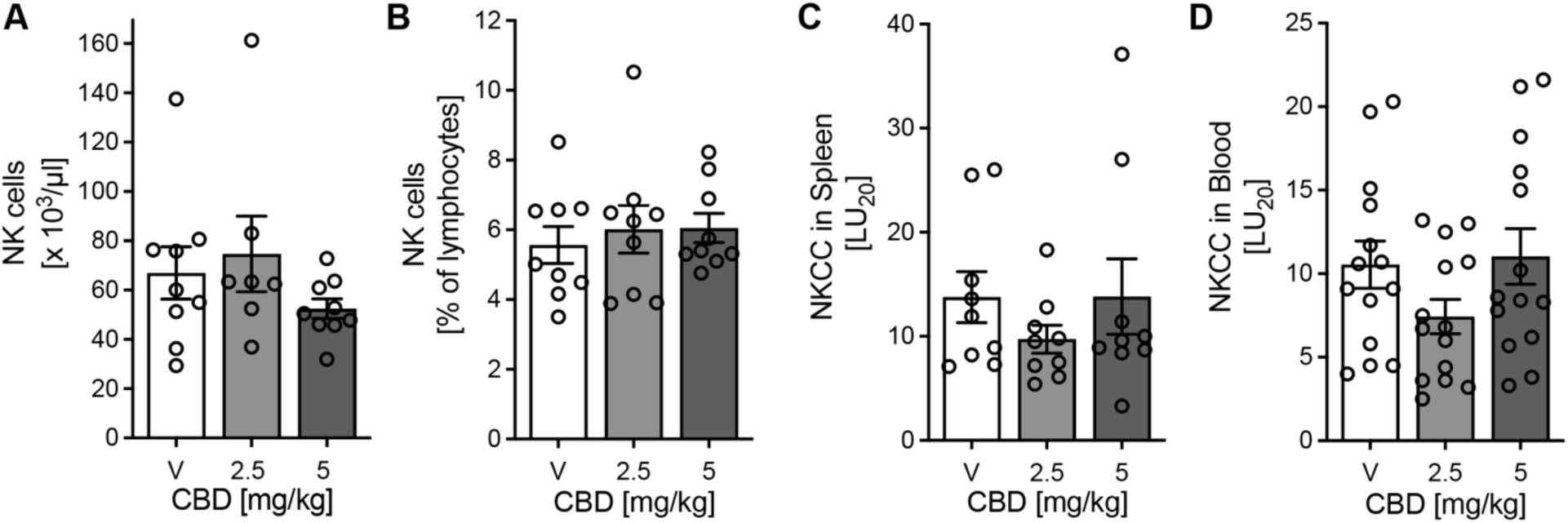
NK cell counts of rats treated with cannabidiol (CBD) for 14 consecutive days. (A) Total NK cell counts. (B) Percentage of total lymphocytes. n = 7-9 per group for A and B. (C) Cytotoxic activity of NK cells towards YAC-1 murine lymphoma tumor target cells in the spleen, n = 9 per group. (D) Cytotoxic activity of NK cells in the blood. n = 9 per group for (C) and n = 14 per group for (D). Data presented as means ± SEM. CBD: cannabidiol, V: vehicle.

CBD administration induced a significant decrease in total spleen weight of 1580 ± 111 mg in the vehicle group, to 1253 ± 77 mg and 1243 ± 77 mg in groups treated with 2.5 and 5 mg/kg/day of CBD, respectively (p = 0.021; F_2,24_ = 4.56, Supp. Fig. 1A). Similarly, spleen weight relative to total bodyweight also decreased from 0.5 ± 0.031 % in the vehicle group to 0.41 ± 0.018 % and 0.41 ± 0.023 % both in animals treated with 2.5 or 5 mg/kg/day of CBD, respectively (p = 0.023; F_2,24_ = 4.46, Supp. Fig. 1B). There were no changes in spleen volume among the groups (p = 0.68, F_2,22_ = 0.39), so significantly reduced spleen density was observed in animals treated with CBD at a dose of 5 mg/kg. Spleen density decreased from 1.079 ± 0.038 μL in the vehicle-treated group to 0.853 ± 0.031 μL in the 5 mg/kg dose-treated group (p = 0.0046, F_2,22_ = 6.925). Additionally, there were no significant differences between blood corticosterone concentrations between the vehicle, 2.5 mg/mL, and 5 mg/mL groups (Supplementary Fig. 4).

### Experiment 2

In rats that received CBD at a dose of 5 mg/kg per day and were pretreated with the vehicle for 14 consecutive days (VC), a significant decrease in the number of splenic leukocytes was observed (Fig. 3A, Two-way ANOVA, p = 0.018, F_1,33_ = 7.119 for effect of CBD). The decrease reached 38.1 % as compared to the group receiving double injections of vehicle (VV) (Two-way ANOVA, Tukey’s post hoc test, VC vs. VV p = 0.007). The same effects were observed in the total lymphocyte number (Supp. Fig. 5). Furthermore, a two-way ANOVA revealed a significant interaction between CBD and AM630 treatment in the case of total numbers of splenic leukocyte (p = 0.021, F_1,33_ = 5.87), lymphocyte (p = 0.020, F_1,33_ = 6.02), B lymphocyte (p = 0.041, F_1,33_ = 4.54), T lymphocyte (p = 0.029, F_1,33_ = 5.22) and T CD8+ lymphocyte (p = 0.011, F_1,33_ = 7.18), but not T CD4+ lymphocyte subset (p = 0.076, F_1,33_ = 3.35).

**Fig. 3.**
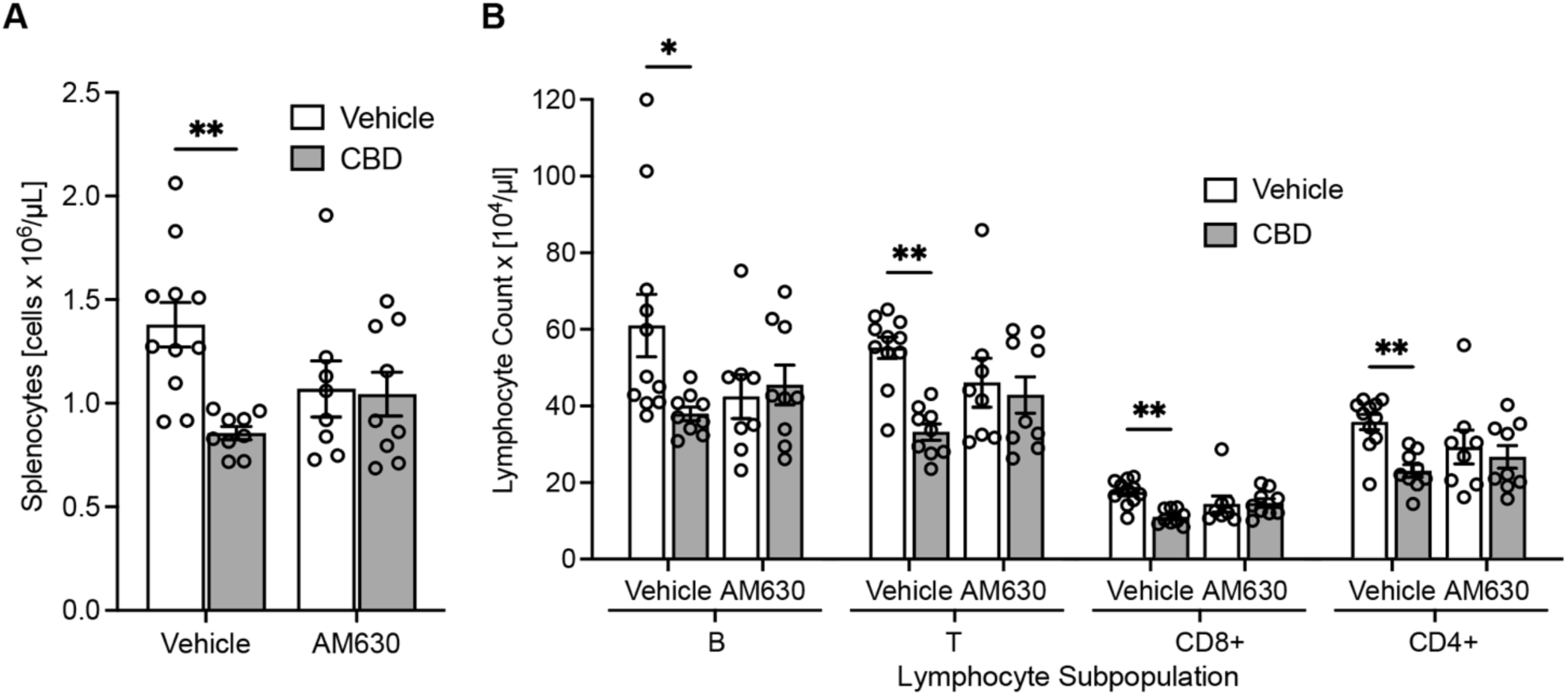
Decreased total leukocyte and lymphocyte number in the spleen produced by administration of cannabidiol (5 mg/kg) for 14 consecutive days is partially blocked by pretreatment with CB_2_ receptor selective antagonist AM630 (1 mg/kg). CB_2_ antagonist by itself also significantly decreased T lymphocyte number. (A) Total splenic leukocyte number, (B) total number of splenocyte subsets: B, T, T CD8+ and T CD4. Two-way ANOVA followed by Tukey’s post hoc test. *p < 0.05, **p < 0.01, ***p < 0.005. Data presented as means ± SEM, n = 8-11 per group. CBD: cannabidiol.

Pretreatment with the CB_2_ antagonist AM630 partially blocked the decrease in leukocyte number produced by CBD administration (AC) as there was no longer a significant decrease in splenocyte number under treatment of both AM630 and CBD (Two-way ANOVA, Tukey’s post hoc test, AC vs. VC p = 0.806). However, animals in the AC group had slightly decreased lymphocyte numbers compared to the VV group similar to the effects produced by AM630 followed by vehicle administration (AV). No significant change in total or percentage NK cell numbers was observed (p = 0.086, F_1,33_ = 2.39, not shown).

There were no significant changes in total lymphocyte numbers between the AV and VV groups (Supp. Fig. 5). However, in terms of the lymphocyte subpopulations, a repetitive tendency for decreased lymphocyte B, T, T CD4+, and CD8+ was observed, but in any case, it did not reach a level of significance. No significant changes in the percentage of lymphocyte subpopulation numbers were observed (data not shown), suggesting a proportional fall in B and T cell numbers. CBD did not affect total or percentage numbers of granulocyte or monocyte numbers in the spleen (data not shown).

CBD administration produced a decrease in total spleen weight (from 890 ± 43 mg in VV to 723 ± 34 mg in VC (Two-way ANOVA, Tukey’s post hoc test, p = 0.002, F_1,33_ = 12.03 for the effect of CBD, Supp. Fig. 6). In terms of relative spleen weight (percentage of total body weight), the effect of CBD administration was significant, (Two-way ANOVA, p = 0.009, F_1,33_ = 7.78 for the effect of CBD), but in the post hoc analysis, the decrease from 0.28 ± 0.01 to 0.25 ± 0.01, for VV and VC, respectively, was not significant (Two-way ANOVA, Tukey’s post hoc test, p = 0.088, Supp. Fig. 6). CBD did not affect spleen volume (p = 0.59, F_1,33_ = 0.296, not shown), and spleen density was reduced by CBD administration, but this tendency was not significant (p = 0.078, F_1,33_ = 3.31, not shown). AM630 pretreatment did not affect spleen weight significantly (Two-way ANOVA, p = 0.479, F_1,33_ = 0.51 for the effect of AM630 pretreatment), but partially prevented the decrease in spleen weight in animals treated with CBD. Furthermore, no significant change was found between VV and AC groups, suggesting blockade of decreased spleen weight by pretreatment with AM630, but no significant interaction between both compounds was revealed (Supp. Fig. 6).

## Discussion

The results of this study revealed that repeated systemic administration of CBD at a dose of 5 mg/kg reduces the total number of lymphocytes and their subsets (B, T, NK, T CD4+, and T CD8+) in the spleen of healthy adult rats. A lower CBD dose of 2.5 mg/kg did not affect spleen leukocyte numbers. NK cells were not involved in this proportional decrease in T and B lymphocytes in the spleen, as their total number, percentage, or cytotoxic activity remained unchanged paralleling previous findings in peripheral blood (Ignatowska-Jankowska et al., 2009). Decrease in not only spleen weight, but also density due to no observed significant difference in volume supports the depletion of lymphocytes in the spleen potentially due to migration. Importantly, the decrease in lymphocytes was partially reversed by pre-treatment with the CB_2_ antagonist AM630. Therefore, there is reason to believe that the lymphopenic effect may be partially mediated *via* CB_2_ receptors and occurs independently of corticosterone levels, suggesting the hypothalamic-pituitary-adrenal (HPA) axis is unlikely to be involved (Besedovsky et al., 1991; Spencer and Deak, 2017).

CBD was found to reduce lymphocyte numbers without an effect on granulocytes and monocytes (Fig. 1, Supp. Fig. 2-3). B and T lymphocytes play an essential role in adaptive immunity, which targets specific antigens rendering it effective against pathogens, and the overactivity of this process may also contribute to autoimmune diseases where host cells are targeted instead of foreign antigens. Meanwhile, granulocytes and monocytes are associated with mechanisms behind innate immunity (Gibbs, 2005; Kobayashi and DeLeo, 2009; Parihar et al., 2010; Shamri et al., 2011). Through its effects exclusively on lymphocytes in this study, CBD may influence adaptive immunity factors specifically, which may be tied to the therapeutic effectiveness of CBD in models of autoimmune diseases such as type I diabetes, RA, or MS, where pathogenic lymphocytes contribute to tissue damage. A question remains of the fate of these lymphocytes that were no longer detected in the spleen. A similar decrease in lymphocyte numbers has been reported in the blood (Ignatowska-Jankowska et al., 2009), and pro-apoptotic effects of CBD on monocytes have been shown to be CB_1_ and CB_2_ receptor-independent (Wu et al., 2018), but less is known about the role of CB_2_ receptors in human lymphocytes (Simard et al., 2022). Induction of apoptosis of splenocytes has been supported by evidence of caspase-8 activation and depletion of intracellular glutathione and thiols in vitro supporting our hypothesis of lymphocyte count decrease due to programmed cell death (Wu et al., 2008; Wu and Jan, 2010).

As an alternative cause of lymphopenia, endocannabinoid 2-arachidonylglycerol (2-AG) has been shown to cause immune cell chemotaxis in a manner that is CB_2_ receptor-dependent, so it is possible that CBD indirectly caused lymphocyte migration, but this could require further examination (Basu et al., 2013; Basu and Dittel, 2011; Cabral and Griffin-Thomas, 2009; Rahaman and Ganguly, 2021). There is evidence of the role of CB_2_ receptors in inhibiting the migration of lymphocytes *in vivo* and *in vitro* (Coopman et al., 2007; Lunn et al., 2006). However, it has been difficult to characterize the role of cannabinoids in their abilities to recruit immune cells as well as interfere with chemoattractants rendering their role complex and context-dependent (Miller and Stella, 2008).

In previous studies, the administration of CBD at a lower dose of 2.5 mg/kg significantly increased the percentage of NK cell numbers in peripheral blood (Ignatowska-Jankowska et al., 2009). This led to the hypothesis that CBD may affect NK cell function. In this study, no effects of CBD on NK cell numbers and NKCC were found suggesting that antitumor effects of CBD may not be associated with increased activity of this cell population. These findings also support the possibility of using CBD as a treatment for patients with cancer or HIV/AIDS, for whom decreased NKCC would be an unwanted effect, with negative consequences for the development of the disease. Furthermore, this highlights the effects CBD as specific to the adaptive immune response without affecting some elements of the innate immune response.

Both endo- and phytocannabinoids are known to directly inhibit immune cell function *via* CB_2_ receptors which are located mainly in the periphery and are very abundant on the immune cells (Buckley et al., 2000; Cencioni et al., 2021; Eisenstein et al., 2007; Galiègue et al., 1995). CBD may affect the immune system through CB_2_ receptors either by direct interaction with receptors or indirectly by affecting endocannabinoid signaling. CBD has been shown to function as a negative allosteric modulator of CB_2_ receptors, as well as an inverse agonist of both CB_1_ and CB_2_ receptors *in vitro* (Martínez-Pinilla et al., 2017; Thomas et al., 2007). Despite its low affinity to CB_2_ receptors, CBD may affect them by prolonging anandamide (AEA) signaling *via* inhibition of fatty acid amide hydrolase (FAAH) which is responsible for the enzymatic breakdown of AEA FAAH (Bisogno et al., 2001; Izzo et al., 2009; Rakhshan et al., 2000). AEA has been shown to have immunosuppressive effects in the spleen such as a decrease in antibody production which was successfully blocked by a CB_2_ receptor antagonist (Eisenstein et al., 2007). Furthermore, the study of CBD in the treatment of inflammatory bowel disease and intestinal inflammation *in vivo* has suggested that the mechanism of FAAH inhibition by CBD is physiologically relevant and promising (Capasso et al., 2008; Pagano et al., 2016). Beyond the scope of this study, the mechanisms behind the role of CB_2_ in distinct affecting specific lymphocyte subpopulation numbers should be further studied *in vivo* to understand their physiological relevance. Additionally, while this study confirms the ability of CBD to induce lymphopenia in the spleen of adult rats, its findings necessitate further validation for its conclusions to be considered for clinical examination or future therapeutic applications.

Regardless of the mechanisms underlying it, CBD-induced lymphopenia may have potential therapeutic benefits in humans. While a decrease in lymphocyte number could impair immune response and worsen prognoses for patients with deficient immune responses, the effects could also be beneficial in the appropriate conditions. It has been shown that depletion in B cells in the periphery may decrease the number of brain lesions and relapse episodes in MS patients (Bar-Or, 2008; Cencioni et al., 2021; Hauser et al., 2008). Therefore, a decrease of B cells in the blood and the spleen could contribute to anti-autoimmune and neuroprotective actions of CBD in MS and possibly other autoimmune diseases.

The present study revealed that repeated CBD treatment decreases lymphocyte numbers not only in the peripheral blood but also in the spleen and CB_2_ receptors are partially involved in its effects *in vivo*. Interestingly, a similar but weaker lymphopenic effect was observed following AM630 administration alone, which suggests that CBD may be acting similarly to a CB_2_ antagonist/inverse agonist, a negative allosteric modulator, or exerting similar effects indirectly. Alternative mechanisms besides CB_2_ may be involved, and further studies are necessary to establish that. Since the lymphopenic effect is observed both in the peripheral blood and spleen, it is possible that it may not be an effect of lymphocyte redistribution but rather due to apoptosis, which should be further examined. Our studies also revealed that the effects of CBD on leukocyte numbers are limited to B and T cells, and the numbers of NK cells and their function were unaffected by CBD administration, suggesting a selective effect of CBD on cells related to adaptive immunity.

## Acknowledgements

This research was supported by Polish Ministry of Science and Higher Education Grants N N303 394036 and N N303 417137 to B. Ignatowska-Jankowska and M. Jankowski. The research was partly financed by the European Union within the European Social Fund in the framework of the project “InnoDoktorant – Scholarships for PhD students, I edition” and the Program of implementing the elements of modern education forms at the University of Gdansk, task: Research grants for PhD students – a chance for the economic development. The project was also supported by the Foundation for Polish Science Ventures Programme (Ventures/2009-4/3), co-financed by the EU European Regional Development Fund. Finally, the authors would like to thank the National Institute on Drug Abuse (NIDA), USA, for drug supply.

## Supplementary Figures

**Supplementary Fig 1.**
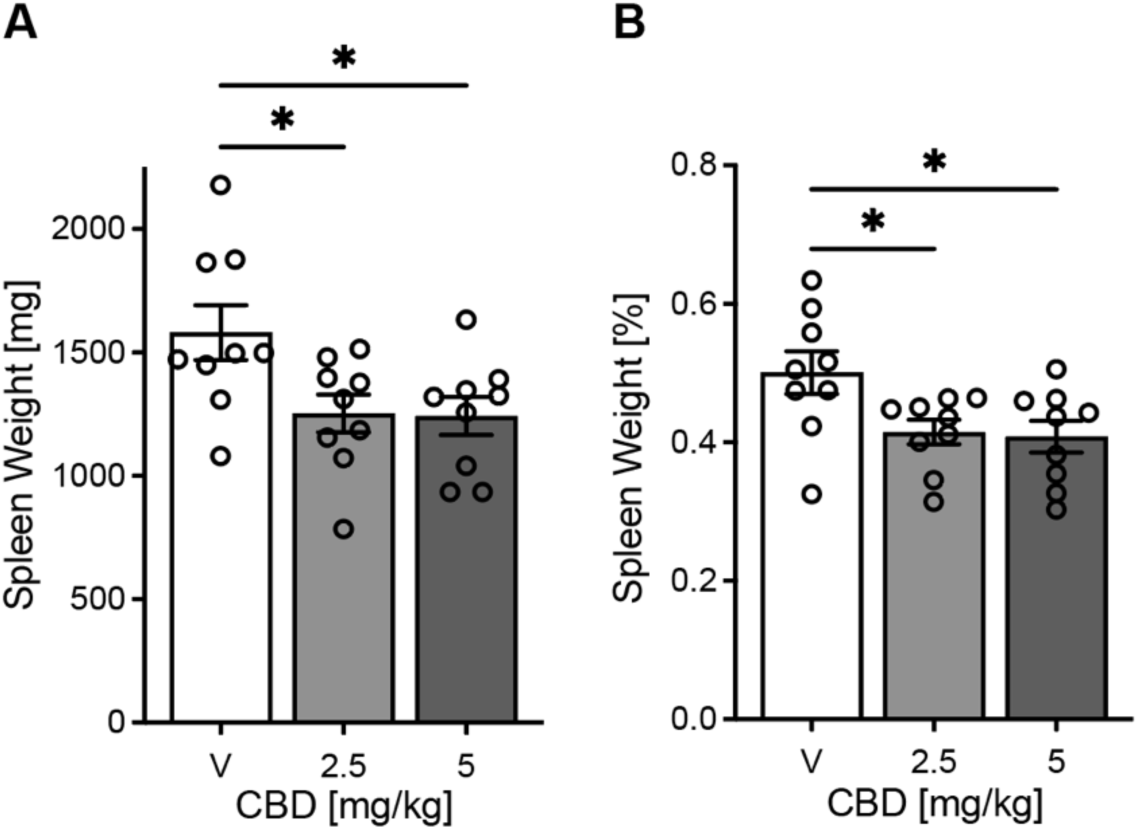
Decreased spleen weight following vehicle, 2.5, and 5 mg/mL cannabidiol administration for 14 consecutive days. (A) Total spleen weight in milligrams (p = 0.021, F_2,24_ = 4.56). (B) Total spleen weight as a percentage of total body weight (p = 0.023, F_2,24_ = 4.46). One-way ANOVA and Dunnett’s post hoc test. n = 27 for A and B. *p < 0.05. CBD: cannabidiol, V: vehicle.

**Supplementary Fig 2.**
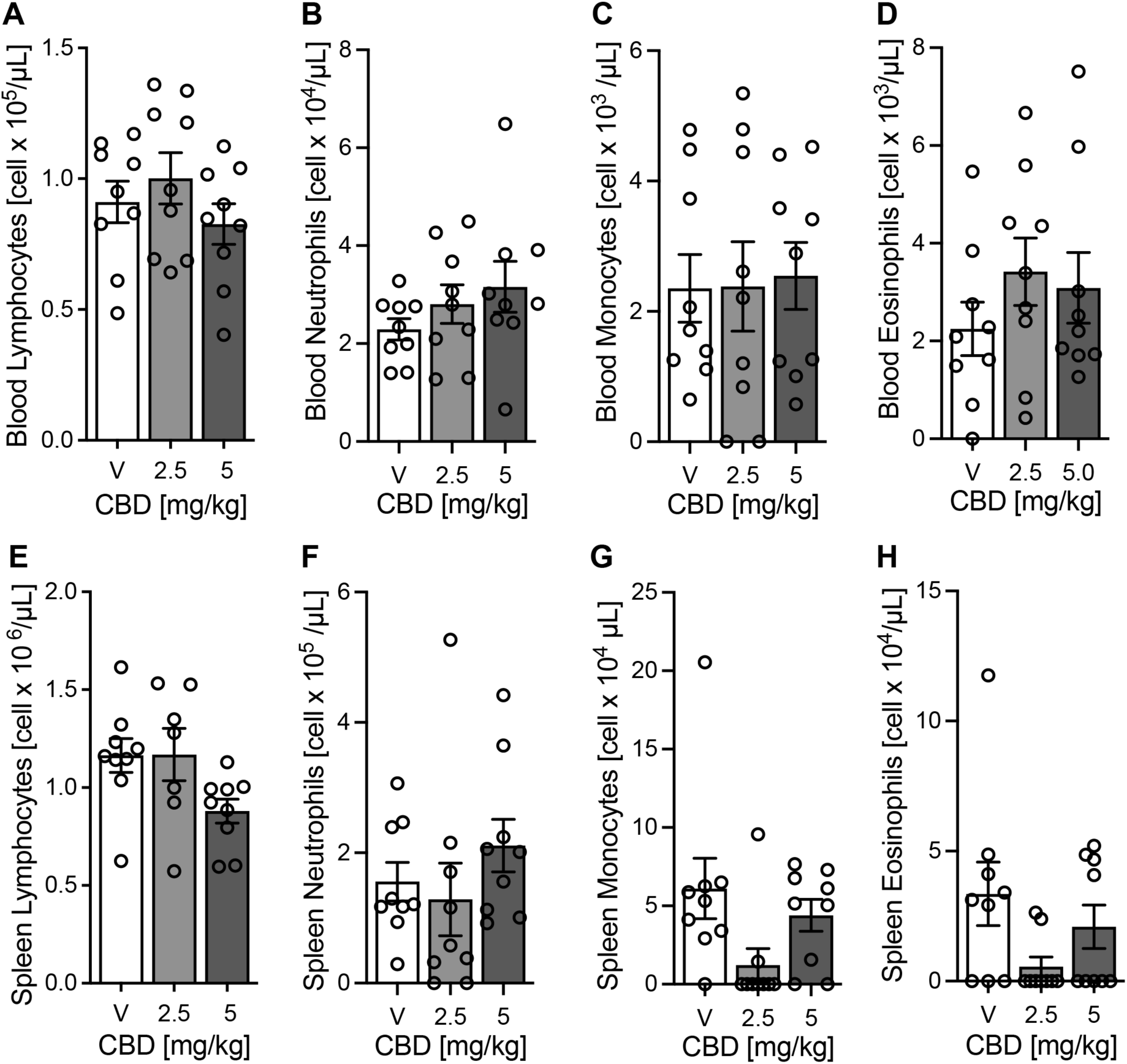
No significant change in the number of granulocytes following CBD administration. Administration of vehicle, 2.5, and 5 mg/mL cannabidiol for 14 consecutive days did not change cell number in (A) lymphocytes, (B) neutrophils, (C) monocytes, and (D) eosinophils in the blood, as well as in the spleen (E-H). Data presented as means ± SEM, n=7-9 per group. CBD: cannabidiol, V: vehicle.

**Supplementary Fig 3.**
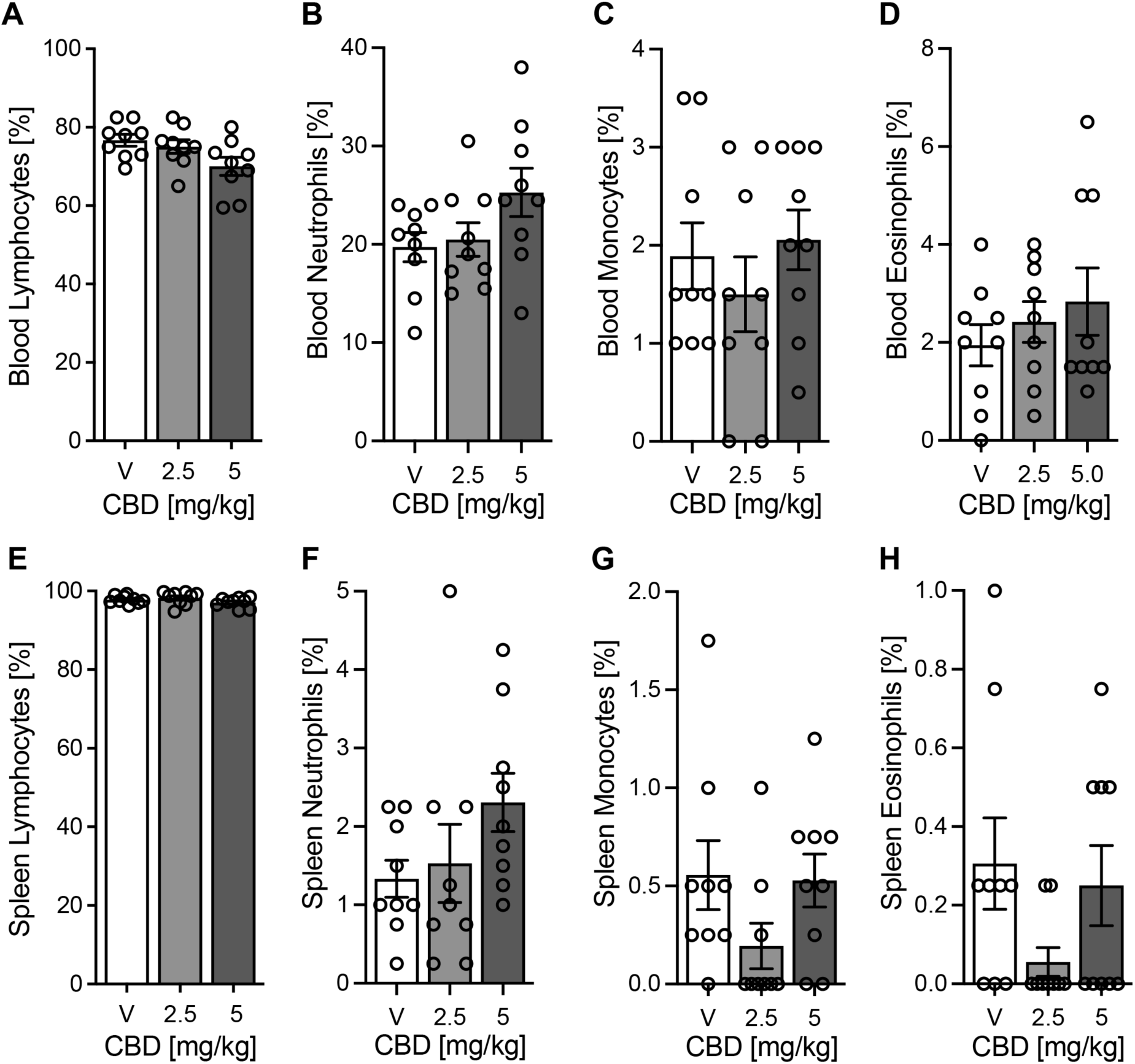
No significant change in the percentage of granulocytes after CBD administration. Administration of vehicle, 2.5, and 5 mg/mL cannabidiol for 14 consecutive days did not change cell number in (A) lymphocytes, (B) neutrophils, (C) monocytes, and (D) eosinophils in the blood, as well as in the spleen (E-H). Data presented as means ± SEM, n=7-9 per group. CBD: cannabidiol, V: vehicle.

**Supplementary Fig. 4:**
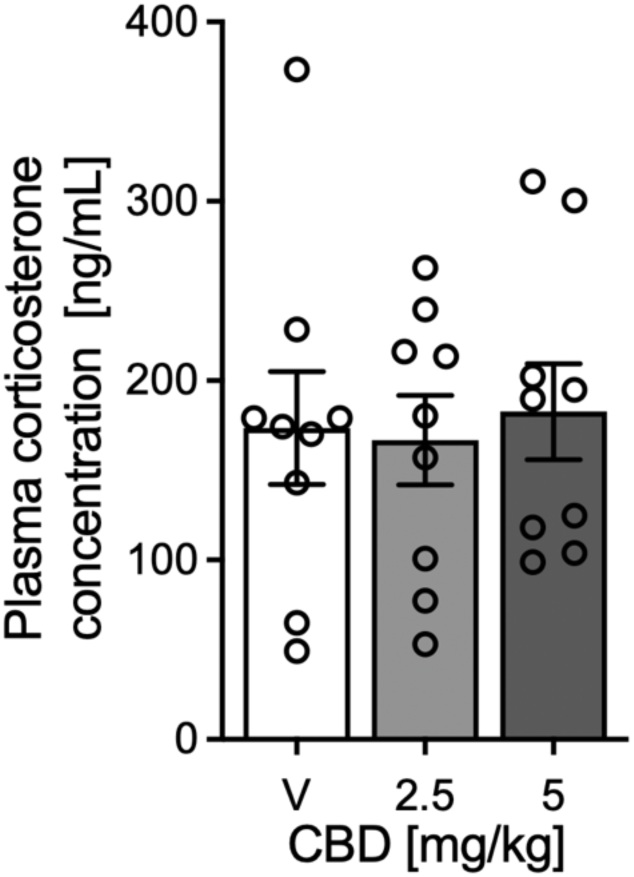
Peripheral plasma corticosterone concentrations in rats injected with vehicle, 2.5, and 5 mg/mL cannabidiol for 14 consecutive days. Data presented as means ± SEM, n=9 per group. CBD: cannabidiol, V: vehicle.

**Supplementary Fig. 5.**
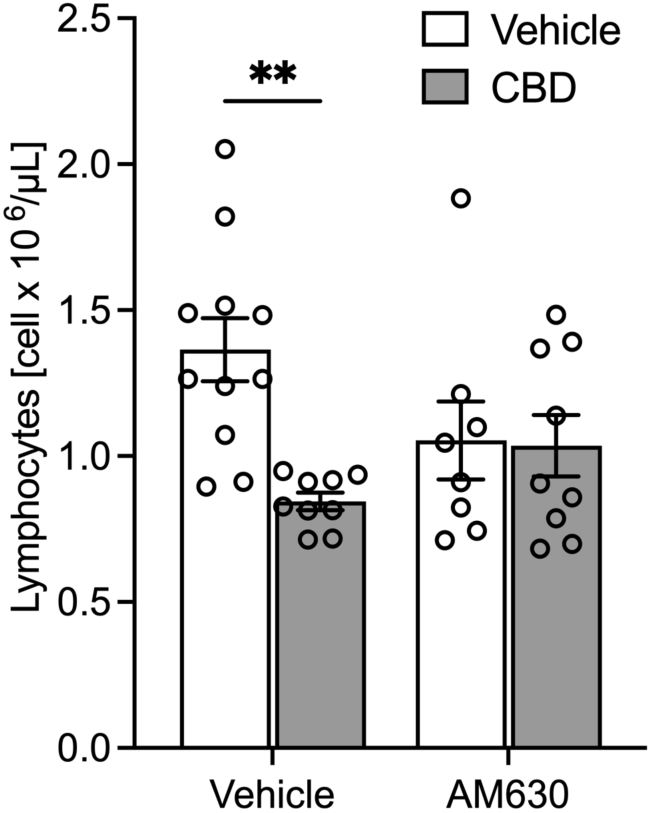
Decreased total lymphocyte number produced by administration of cannabidiol (5 mg/kg) for 14 consecutive days is partially blocked by pretreatment with CB_2_ receptor selective antagonist AM630 (1 mg/kg). Two-way ANOVA followed by Tukey’s post hoc test. *p < 0.05, **p < 0.01, ***p < 0.005. Data presented as means ± SEM, n=8-11 per group. CBD: cannabidiol.

**Supplementary Fig. 6.**
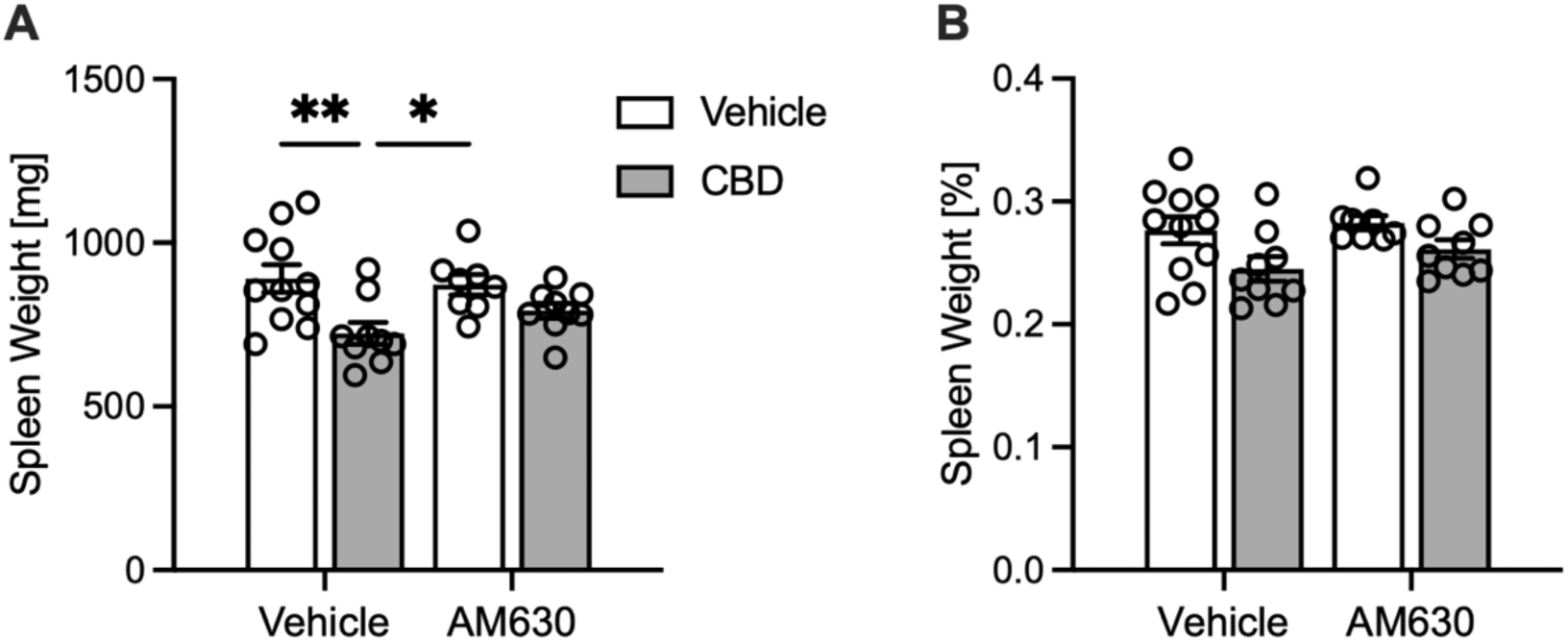
Change in spleen weight following administration of cannabidiol (5 mg/kg) for 14 consecutive days is partially blocked by pretreatment with CB_2_ receptor selective antagonist AM630 (1 mg/kg). (A) Total spleen weight in milligrams, (B) Total spleen weight as a percentage of total body weight. Two-way ANOVA and Tukey’s post hoc test. Data presented as means ± SEM=8-11 per group. CBD: cannabidiol.

